# Evolution of life cycles and reproductive traits: insights from the brown algae

**DOI:** 10.1101/530477

**Authors:** Svenja Heesch, Martha Serrano-Serrano, Rémy Luthringer, Akira F. Peters, Christophe Destombe, J. Mark Cock, Myriam Valero, Denis Roze, Nicolas Salamin, Susana Coelho

## Abstract

Brown algae are characterized by a remarkable diversity of life cycles, sexual systems, and reproductive modes, and these traits seem to be very labile across the whole group. This diversity makes them ideal models to test existing theories on the evolution of alternation between generations, and to examine correlations between life cycle and reproductive life history traits. In this study, we investigate the dynamics of trait evolution for four life-history traits: life cycle, sexual system, level of gamete dimorphism and gamete parthenogenetic capacity. We assign states to up to 70 species in a multi-gene phylogeny of brown algae, and use maximum likelihood and Bayesian analyses of correlated evolution, taking phylogeny into account, to test for correlations between life history traits and sexual systems, and to investigate the sequence of trait acquisition. Our analyses are consistent with the prediction that diploid growth may evolve because it allows the complementation of deleterious mutations, and that haploid sex determination is ancestral in relation to diploid sex determination. However, the idea that increased zygotic and diploid growth is associated with increased sexual dimorphism is not supported by our analysis. Finally, it appears that in the brown algae isogamous species evolved from anisogamous ancestors.

## INTRODUCTION

The life cycle of an organism is one of its most fundamental features and influences the evolution of a variety of traits, including mode of reproduction, developmental processes, mode of dispersal, adaptation to local environment and ecological success. A wide variety of different life cycles are found within eukaryotes, and one of the great challenges of evolutionary biology is to understand how this diversity has evolved, and how each type of life cycle is stably maintained on an evolutionary timescale (Valero et al. 1992; Mable and Otto 1998; Otto and Gerstein 2008; Cock et al. 2014).

The sexual life cycle of eukaryotes involves the fusion of two gametes to form a zygote, followed by meiosis. Such life cycles can be divided into three main types (Bell 1982; Valero et al. 1992; Coelho et al. 2007; Otto and Gerstein 2008). In organisms with haplontic cycles, mitosis only occurs during the haploid phase of the life cycle, and syngamy is followed immediately by zygotic meiosis, without any mitotic division of the zygote. In diplontic life cycles, mitosis occurs exclusively in the diploid phase, and meiosis takes place immediately before gamete formation. Between these two ‘extremes’ lie diplohaplontic (or haploid-diploid) life cycles, in which mitosis occurs both during the haploid and diploid phases. Both phases of the cycle may remain unicellular (e.g. budding yeast) or develop into multicellular organisms during either one phase (e.g. some fungi genera like *Ustilago*) or during both phases (brown, green, red macroalgae). In photosynthetic organisms, multicellular haploid phases are usually termed gametophytes, with gametes formed in specific organs called gametangia. Diploid mitosis leads to asexual reproduction in unicellular lineages (e.g. diatoms) and to somatic growth and differentiation in multicellular organisms such as vascular plants. In photosynthetic organisms, multicellular diploid phases are called sporophytes, and haploid meiospores are produced via meiosis. Diplohaplontic life cycles may be iso-or heteromorphic. For the latter, the dominant phase may be haploid (such as in mosses) or diploid (such as in vascular plants and kelps). Asymmetry in terms of the length and complexity of the haploid and diploid phases can be very strong and can eventually lead to transitions towards diplontic or haplontic life cycles.

The structure of an organism’s life cycle also has important consequence for the evolution of its sex determination system (Coelho et al. 2018). Haploid sex determination is common in diplohaplontic lineages such as in brown algae, where either a single gametophyte produces gametes of both sexes (a monoicous system, Table 1), or male and female gametes are produced by different haploid gametophytes (dioicous systems, e.g. in mosses and kelps). In gymnosperms and angiosperms (which also have diplohaplontic life cycles), sex is determined in the diploid phase (since sexual differentiation occurs when male and female reproductive organs develop), and the organism may be co-sexual (monoecious or hermaphroditic) if a single sporophyte produces female and male gametes (eggs and sperm) or dioecious if male and female gametes are produced by two different individuals. Correlations between the type of sexual system and life history features such as spore size, antheridium number, ploidy level and diversification rate are relatively well studied in angiosperms and mosses (Villarreal and Renner 2013; Goldberg et al. 2017) but studies of other eukaryotic groups are virtually inexistent.

**Table 1.** Description of the traits studied, categories and discrete states. Note that some of the discrete traits were also treated as continuous traits (male gamete size for instance).

|  | Category | States | Description |
| --- | --- | --- | --- |
| Life cycle | Haplo-diplontic | Haplo-diplontic haploid dominant<sup>1</sup> | Life cycle with both haploid and diploid mitosis, with dominant gametophyte haploid generation |
|  |  | Haplo-diplontic haploid = diploid | Life cycle with both haploid and diploid mitosis, with equal dominance of gametophyte and sporophyte generations |
|  |  | Haplo-diplontic diploid dominant<sup>1</sup> | Life cycle with both haploid and diploid mitosis, with dominant sporophyte (diploid) generation |
|  | Diplontic | Diplontic | Life cycle with no haploid mitosis, the haploid phase is limited to gametes |
| Sexual system | Haploid sex determination | Monoicous | Haploid phase sex determination, where both gamete types are produced by the same haploid gametophyte. |
|  |  | Dioicous | Haploid phase sex or mating type determination, with genetically distinct gametophytes corresponding to each sex |
|  | Diploid sex determination | Dioecious | Diploid phase sex-determination, with genetically distinct sporophytes corresponding to each sex |
|  |  | Monoecious | Diploid phase sex determination, where both male and female organs are produced by the same diploid sporophyte |
| Gamete size | Female gamete with flagella | Isogamous<sup>2</sup> | Male and female gametes with small size difference (but different behaviour/physiology) |
|  |  | Anisogamous | Male and female gametes of clearly different size, both with flagella |
|  | Female gamete without flagella | Oogamous | Female gamete much larger and lacking a flagellum |
| Parthenogenesis | No parthenogenesis | No parthenogenesis | No parthenogenesis capacity in either gamete |
|  | Parthenogenesis | Female gametes only | Only female gametes capable of parthenogenesis |
|  |  | Male and female gametes | Male and female gametes capable of parthenogenesis |

<sup>1</sup>The term "dominant" is defined here as the generation that presents larger size and higher complexity in terms of morphology (number of different cell types, number of tissues and organs)
<sup>2</sup>For simplicity, we code as "isogamous" algae that have almost imperceptible size differences between male and female gametes, but note that in the brown algae there is always an asymmetry (at least in terms of physiology and behaviour) between male and female gametes.

One important feature of sexual life cycles in eukaryotes is the degree of similarity between male and female gametes. This ‘gamete dimorphism’ is a continuous trait, and a number of models have been proposed to explain how anisogamous organisms could evolve from an isogamous ancestor (Charlesworth 1978; Hoekstra 1980; Randerson and Hurst 2001). The evolution of anisogamy establishes the fundamental basis for maleness and femaleness, and leads to an asymmetry in resource allocation to the offspring, leading in many cases to sexual selection (Billiard et al. 2011). Anisogamy and oogamy have arisen repeatedly across the eukaryotes, and these systems are thought to be derived from simpler isogamous mating systems, either due to disruptive selection generated by a trade-off between the number of offspring produced and offspring survival (e.g., Charlesworth 1978; Parker 1978; Bell 1982; Bulmer and Parker 2002), to selection to maximize the rate of gamete encounter (e.g., Cox and Sethian 1984; Hoekstra 1987; Dusenbery 2000; Togashi et al. 2012) or as a mechanism to reduce cytoplasmic conflicts (e.g., Hurst & Hamilton1 1992, Hutson & Law 1993, Hurst 1995).

Differences in gamete size in anisogamous and oogamous species may influence other reproductive characteristics. In particular, gamete size may be one of the factors that determines whether a gamete is capable of undergoing asexual reproduction through parthenogenesis, should it fail to encounter a gamete of the opposite sex (Hoekstra 1980, Billiard et al 2011). Parthenogenesis is a form of asexual reproduction, in which gametes develop in the absence of fertilisation, and is commonly found in land plants, algae and invertebrate organisms, as well as in a number of vertebrate species (e.g. Dawley and Bogart 1989). In animals and land plants, parthenogenesis has been mostly described for females only, but in organisms with moderate levels of gamete dimorphism such as some brown algae, development from both male and female gametes in the absence of fertilisation is quite common, at least under laboratory conditions (e.g. Oppliger et al. 2007; Bothwell et al. 2010).

The different types of life cycles have evolved independently and repeatedly in different eukaryotic groups, and this is also the case for the types of sexual systems. Testing evolutionary hypotheses regarding the causes and consequences of life history trait diversity requires data from multiple species placed in a phylogenetic context. Such comparative studies have been hampered by a lack of accessible data regarding life cycles, sexual systems and sex determination mechanisms across the eukaryotic tree of life, and most specifically in groups outside animals and land plants. While knowledge has been recently growing on the green lineage with studies extending to bryophytes and volvocine algae (Villarreal and Renner 2013; Hanschen et al. 2018), we still lack views on other eukaryotic groups, that should help us understand the general principles underlying the evolution of these traits.

The brown algae represent a fascinating group for studies of the evolution of life cycles and reproductive traits (Bell 1997; Clayton 2009). Brown algae exhibit a remarkable range of life cycle and sexual traits. Most brown algae have diplohaplontic life cycles, in which a haploid gametophytic generation alternates with a diploid sporophytic generation (Figure 1). These two generations show variable levels of dimorphism, ranging from isomorphic (with gametophytes and sporophytes of the same size and appearance) to strongly heteromorphic (where the generations are of different size and/or appearance) with either a dominant gametophytic or a dominant sporophytic generation. In two orders, the Fucales and the Tilopteridales, diplontic life cycles have evolved, with all members of the Fucales being diplonts, but only a single species in the Tilopteridales. In diplonts, the gametophytic generation is reduced to a single cell (the gamete), male and female gametes being released directly from the diploid sporophyte, either from the same (monoecy/hermaphroditism) or from two separate thalli (dioecy). In contrast, in all other sexual brown algal species, sexuality is expressed in the haploid phase (gametophyte), with male and female gametes either produced on the same thallus (monoicy) or on two separate male and female gametophytes (dioicy) (Silberfeld et al. 2010; Luthringer et al. 2015). Here, we will use here the term ‘monoecy’ to describe Fucales species that are co-sexual, although the term hermaphrodite is also employed in the literature, because male and female structures are produced in the same conceptacle.

**Figure 1.**
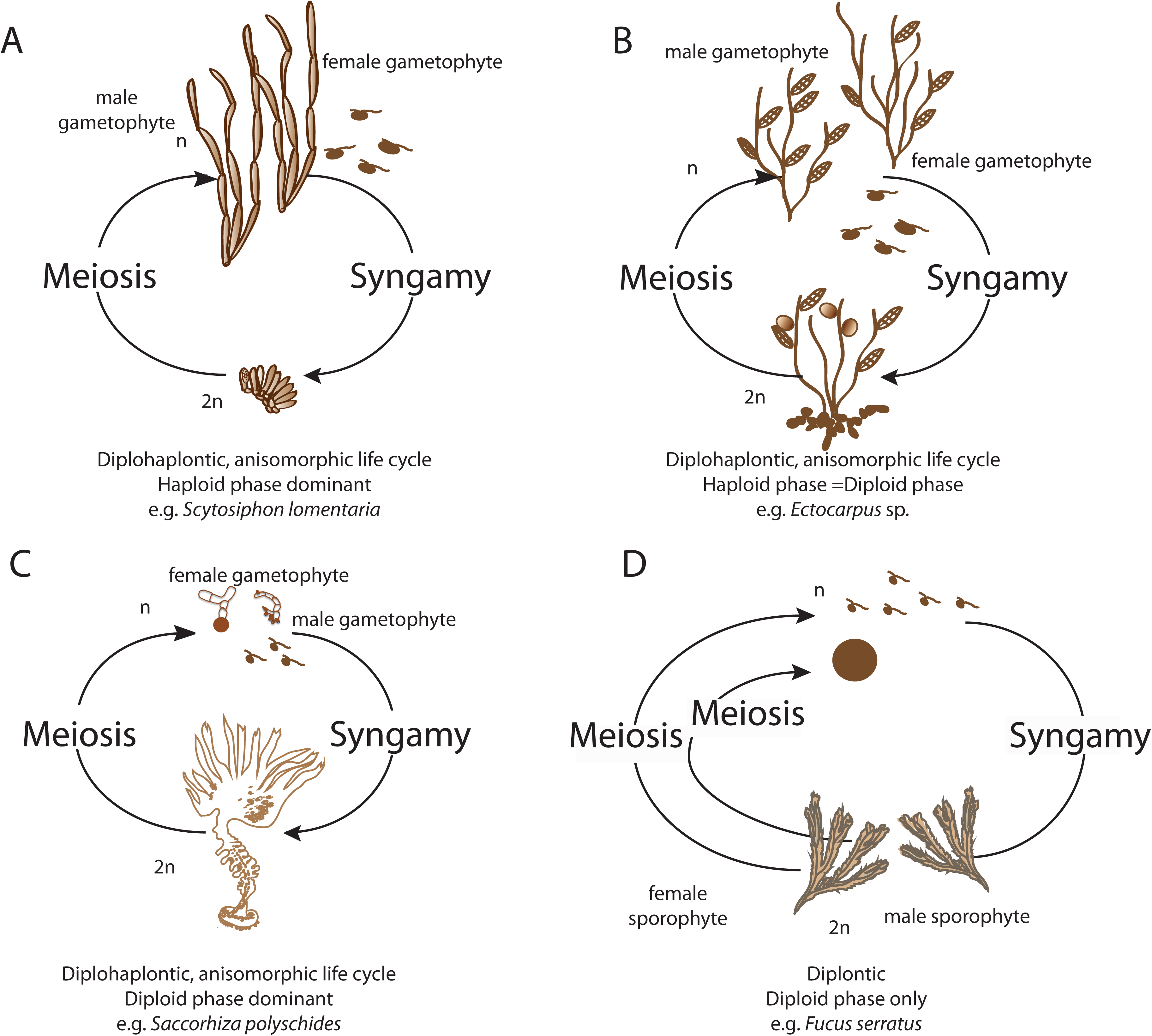
Schematic illustration of sexual life cycles of representative brown algae. (A) *Scytosiphon lomentaria* (Lyngbye) Link: diplohaplontic, heteromorphic life cycle, haploid dominant; near-isogametes (B) *Ectocarpus* sp. (Dillwyn) Lyngbye: diplohaplontic, isomorphic life cycle, with similar dominance in haploid and diploid phases; near-isogametes; (C) *Saccorhiza polyschides* (Lightfoot) Batters: diplohaplontic, heteromorphic life cycle, with diploid dominance (D>>H); oogamous; (D) *Fucus serratus* L.: diplontic life cycle, only diploid phase; oogamous. H=haploid phase; D=diploid phase.

In 1997, Bell used the diversity of life cycles within the brown algae to test hypotheses on the evolution of life cycles; in particular, whether evolution generally proceeds towards an increase of the diploid phase at the expense of the haploid phase (Clayton, 1988), and whether a positive association between a prolonged haploid phase and the rate of inbreeding (as predicted by theories based on the effect of deleterious alleles, Otto and Marks 1996) is observed (using gametophyte monoicy as a proxy for inbreeding by assuming that gametophytic selfing may occur). However, his study was based on a phylogenetic tree including only 14 species, and evolutionary relationships between brown algal orders were poorly resolved, making it difficult to test his assumptions.

In this study, we have exploited a well-resolved phylogeny of 70 species of brown algae (Silberfeld et al. 2010) to understand how life cycle and reproductive traits evolve across the brown algae. We performed an extensive literature review to recover information for life cycle and reproductive traits across the brown algae (Supplemental Dataset). We estimated ancestral states for each of the traits, as well as the number of transitions between states and their relative timing, and assessed correlations between the life cycle and reproductive traits. These analyses have allowed us to describe the evolution of life cycles and reproductive traits across the brown algal phylogeny, and to assess a number of long standing hypotheses about the evolution of life cycle and reproductive traits such as: 1) the possibility that diploid growth evolved because it allows the complementation of deleterious mutations (Crow and Kimura 1978, Perrot et al. 1994, Jenkins and Kirkpatrick 1995, Otto & Goldstein 1992, Otto & Marks 1996), 2) that increased zygotic and diploid growth is associated with increased sexual dimorphism (Bell, 1994), 3) that haploid sex determination is ancestral in relation to diploid sex determination, 4) that anisogamous species evolved from isogamous ancestors (Parker et al 1972, Bell 1978, Charlesworth 1978). We also test additional hypotheses, including the possibility that gamete size influences the capacity for asexual reproduction through parthenogenesis (Luthringer et al. 2015), and we discuss the macro-evolutionary dynamics of transitions between sexual systems in the brown algae.

## RESULTS

### Ancestral state estimations and transitions between states

We used sequence data from 131 brown algae species to infer a phylogenetic tree (Figure 2). We estimated the ancestral state of each of the four main sexual traits: type of life cycle, type of sexual system, level of gamete dimorphism, and parthenogenetic capacity. Definitions of the life cycle and sexual terms used in this study are provided in Table 1. Our ancestral state reconstructions inferred equal rates of transition (ER model) between states for all traits, except for the trait ‘sexual system’ where rates were different between states but symmetrical (SYM model gain or loss of a trait). These patterns indicate an overall complex evolutionary history for all sexual traits, involving multiple gains and losses (Figure 3, Table 2).

**Table 2.** Analysis of the ancestral states for each studied trait.

| Trait | Description | No. species | Best Model | Mean rates (SD)<sup>1</sup> | Root state (prob)<sup>2</sup> | Mean No. transitions (SD) | Min time of transition (60% recons, and SD)<sup>3</sup> |
| --- | --- | --- | --- | --- | --- | --- | --- |
| Life cycle | 0: Hap>>Dip, 1: Hap=Dip, 2: Hap<<Dip, 3: Dip | 68 | ER | 0.00135 (1.1e-4) | 0: 0.00117, 1: 0.39560, 2: 0.56764, 3: 0.03559 | 1→0: 1 (0.6), 1→2: 1 (0.9), 1→3 :1 (0.7), 2→0: 1 (0.7), 2→1: 5 (1.5), 2→3: 2 (0.8) | 2→0: 57.5 My (5.05), 2→1: 81.5 My (23.55), 2→3: 74.5 My (21.41) |
| Sexual system | 0: monoicous, 1: dioicous, 2: monoecious, 3: dioecious | 66 | SYM | 0>1: 0.00515 (4.7e-4), 0>2: 0.00000 (4.1e-6), 0>3: 0.00057 (5.1e-4), 1>2: 0.00102 (1.3e-4), 1>3: 0.00000 (0.000), 2>3: 0.01180 (1.2e-3) | 0: 0.02361, 1: 0.96346, 2: 0.01228, 3: 0.00065 | 1→0: 8 (0.9), 1→2: 2 (0.4), 1→3 :1 (0.2), 2→0: 1 (0.2), 2→3: 3 (1.8), 3→2: 2 (1.3) | 1->0: 110.5 My (30.02), 1->2: 74.5 My (21.79), 2->3: 17.5 My (5.34), 3->2: 25.5 My (7.65) |
| Gamete size | 0: isogamy, 1: anisogamy, 2: oogamy | 69 | ER | 0.00236 (1.8e-04) | 0:0.00118, 1:0.00029, 2:0.99854 | 0→1: 2 (0.8), 1→0: 3 (0.8), 2→0: 2 (0.8), 2→1: 8 (1.1) | 0->1: 56.5 My (16.6), 2->0: 113.5 My (33.05), 2->1: 87.5 My (25.28) |
| Parthenogenesis | 0: Nobody, 1: only female, 2: female and male | 40 | ER | 0.00288 (2.4e-4) | 0: 0.59786, 1: 0.16131, 2: 0.24082 | 0->1: 1 (1.2), 0→2: 2 (1.4), 1→0: 2 (1.4), 1→2: 4 (1.9), 2→1: 1 (1.8) | 0->1: 26.5 My (7.94), 0->2: 43.5 My (14.86), 1->2: 85.5 My (24.97), 2->1: 34.5 My (10.25) |

<sup>1</sup>Rates are the mean and sd values of the optimization in a sample of 100 posterior trees. <sup>2</sup>Probability of the root is the mean over 100 reconstructions <sup>3</sup>Stochastic mapping analysis and time where 60% of the reconstructions that have at least 1 transition ER: a single transition rate between all states SYM: forward and reverse transitions are the same ARD: each rate is a unique parameter

**Figure 2.**
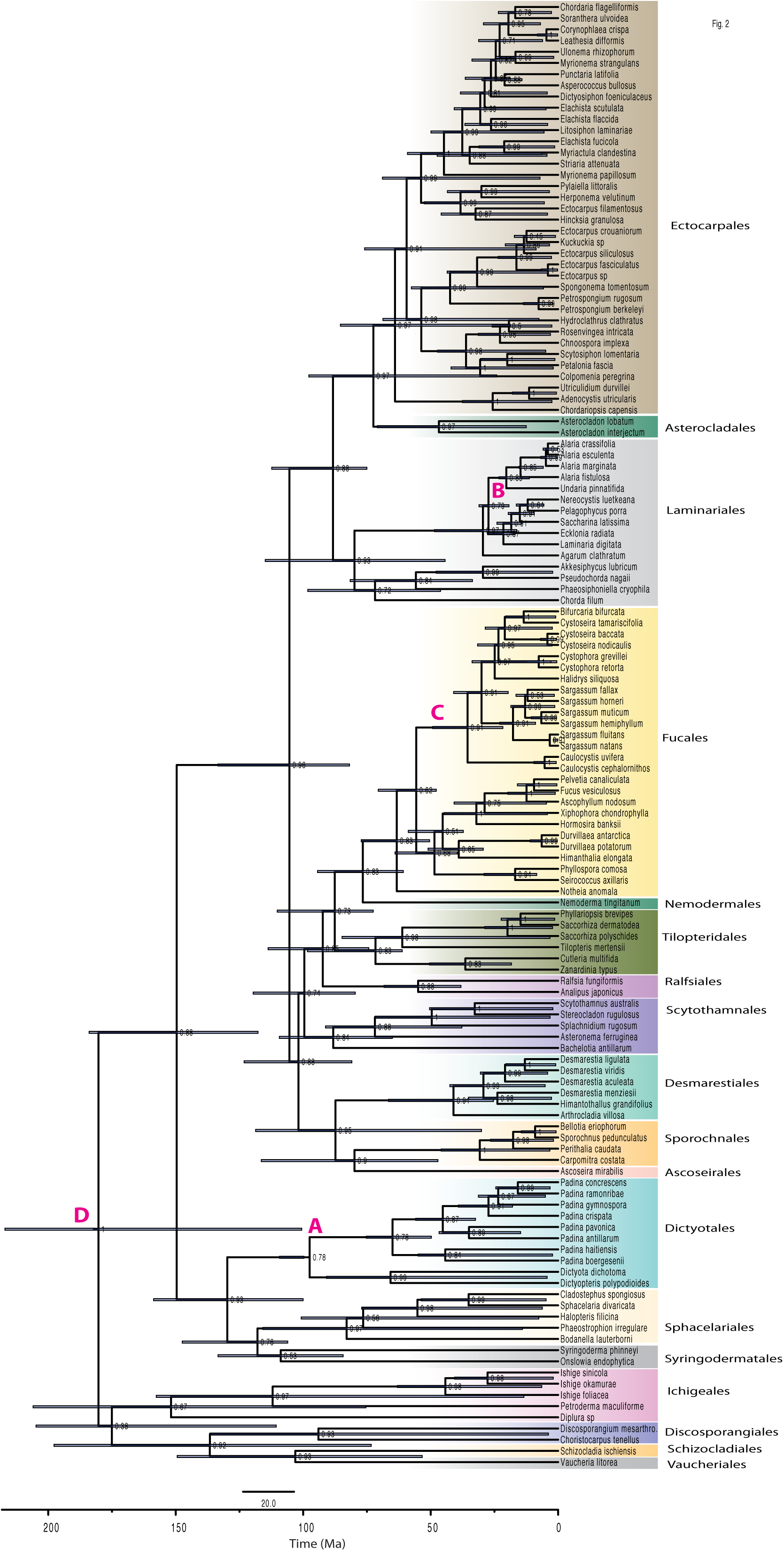
Phylogenetic tree using Bayesian analyses in BEAST. Node numbers indicate the posterior Bayesian support, node bars represent the 95% HPD (Highest Posterior Density) for the divergence times.

**Figure 3.**
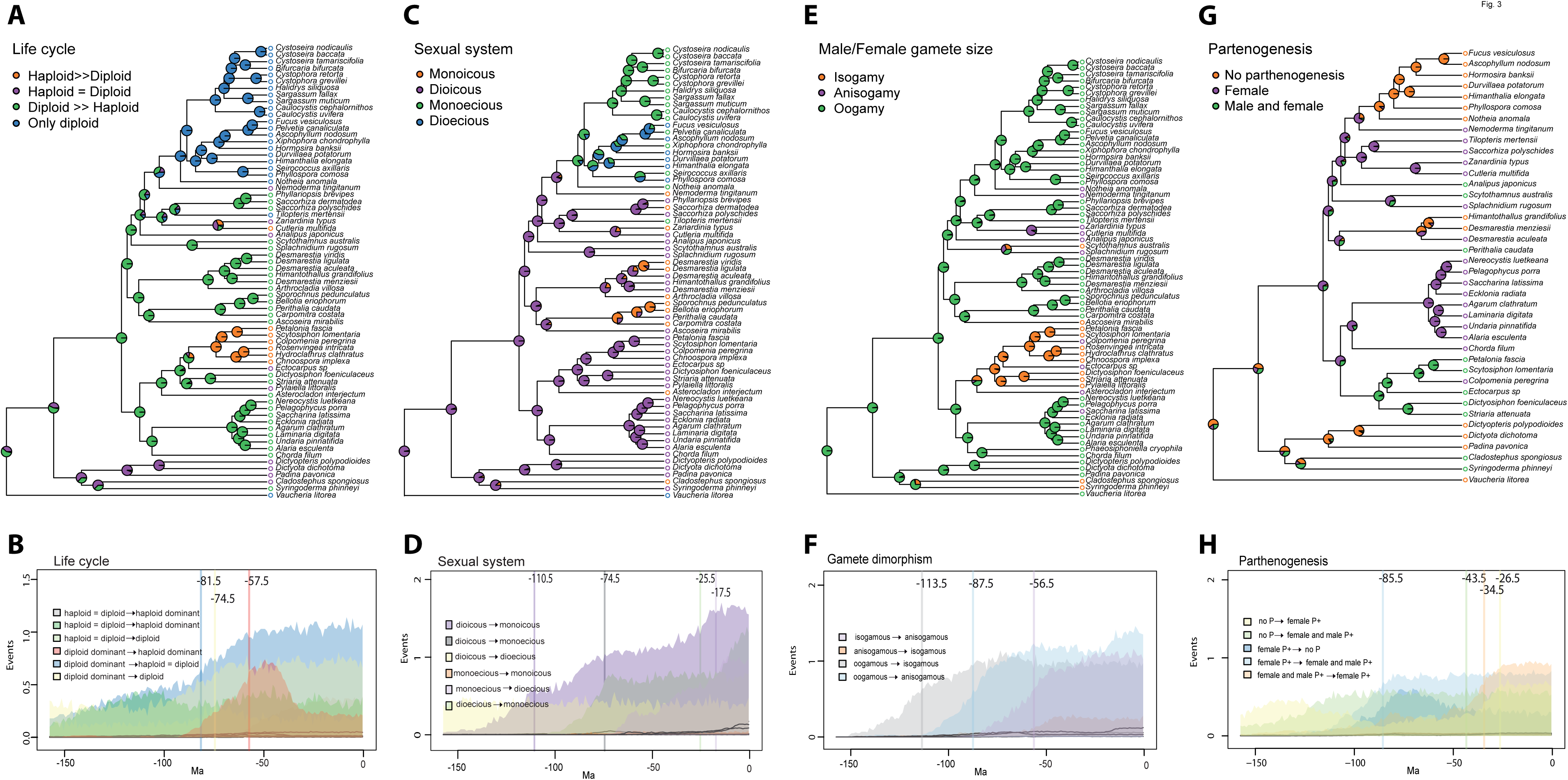
Maximum likelihood ancestral state reconstructions for the four-brown algal life-history traits. Pie charts and colours at each node represent the probabilities for each state. Colours at the tips represent the species states (A, C, E, F). Estimated number of transitions through time for the corresponding four life-history traits (B, D, F, H). Coloured densities identify the mean number of events for each possible transition. Vertical lines and numbers denote the minimum age as the point in time where at least one transition is recorded in 60% of the reconstructions. A and B) Male and female gamete size; C and D) Sexual system, E and F) Type of life cycle, G and H) Parthenogenetic capacity.

#### Life cycle

On the basis of ancestral state reconstructions, the ancestor of all brown algae had a diplohaplontic life cycle, with either isomorphic multicellular generations or with a larger and morphologically more complex diploid than haploid generation (Figure 3A, Table 2). Transitions between life cycles occurred most frequently from diploid-dominant to equally dominant generations (i.e., involving a decrease in complexity of the sporophyte and concomitant increase in the complexity of the gametophyte; Figure 3A). A change of dominance from a diploid-dominant to a haploid-dominant life cycle occurred for the first time at least 57.5 (±5.05) My ago (Figure 3A-B). Transitions from a diploid-dominant to a fully diploid life cycle occurred twice, about 74.5 (±21.41) My ago, in the ancestor of *Tilopteris mertensii* (Turner) Kützing and in the ancestor of the diploid order Fucales. Note however that *Tilopteris mertensii* is a rather particular case within Tilopteridales (Kuhlenkamp and Müller 1985) and emergence of monoecy in this species should be interpreted with caution.

Overall, our analysis indicated that the dominance relationship between life cycle generations has been a labile trait in the brown algae, the diplontic life cycle being the only irreversible state.

#### Sexual system

The ancestral brown alga is predicted to have exhibited haploid sex determination and was most likely dioicous (separate sexes during the haploid phase) (Figure 3A-D, Table 2), but several independent transitions towards monoicy have occurred (Figure 3C-D). The transition from haploid to diploid sex determination, which involved a transition from dioicy (separate sexes on different haploid gametophytes) to monoecy (both sexes on the same diploid sporophyte), occurred about 74.5 My ago. This transition was simultaneous with the transition from a diplohaplontic to a diplontic life cycle (Figure 3B, 3D). Separate sexes during the diploid stage of the life cycle (dioecy) emerged more recently, around 17.5 Mya, in families of the order Fucales, with the exception of the Sargassaceae and Notheiaceae, which remained monoecious. Further transitions back to monoecy occurred in several genera of the Fucaceae (Xiphophora, Pelvetia and Fucus) (Table 2, Figure 3C-D).

Overall, our analysis suggests that the transition to diploid sex determination is irreversible and concomitant with a change in type life cycle (from diplohaplontic to diplontic life cycle). In contrast, transitions between separate sexes and combined sexes occurred frequently, either in the haploid or in the diploid phase.

#### Sexual dimorphism

Regarding gamete size dimorphism, our analysis suggests that oogamy is most likely the ancestral state in the brown algae (Table 2, Figure 3E-F). The oldest transition took place around 114 My ago, from oogamy to isogamy in the lineage leading to the basal brown algal orders Sphacelariales and Syringodermatales. Another independent transition from oogamy to isogamy took place in the Ascoseirales. The Scytothamnales include both isogamous and anisogamous taxa so the direction of the transition is unclear. Transitions from oogamy to anisogamy are the most frequent transition. Taken together, the results indicate that gamete size dimorphism level is a remarkably labile trait in the brown algae.

#### Parthenogenesis

The gametes of the ancestral brown algae are predicted to have been unable to perform asexual reproduction through parthenogenesis (Figure 3G-H). The initial transition from absence of parthenogenesis to female gamete parthenogenesis could not be accurately traced in time along the early diverging branch separating the subclass Fucophycideae from the earlier branching Dictyophycidae. The length of this branch renders identification of the transition during 1 My time bins impossible, as most events fall in different time periods and agreement between reconstructions is very low. The oldest traceable transition that could be timed, at around 85.5 My, and also the one with the highest frequency, was from female-only parthenogenesis to parthenogenesis of both female and male gametes. Note that parthenogenesis is the trait with the lowest sampling, as there is very limited data about this trait in the literature.

### Generation dominance and sexual system

Transitions towards the dominance of haploid phase were found to be more frequent when the sexual system was monoicous (Figure 4, q21>>q43), and overall, switches in life cycle phase dominance probabilities were higher in monoicous compared to dioecious species, whatever the direction (q21 and q12 >q43 and q34; Table 4, Figure 4, Figure S1). In other words, monoicous species exhibit a higher turnover in terms of generation dominance. Moreover, transitions from monoicous to dioicous states were slightly more frequently observed than transitions from dioicous to monoicous, regardless of life cycle phase dominance (Bayes Factor of 3.51 in favour of the dependent model with q24 ∼ q13 > q42 ∼ q31, Figure 4, Figure S1).

**Table 3.** Analysis of correlations between traits.

| *Continuous versus discrete* |  |  |  |  |  |  |  |
| --- | --- | --- | --- | --- | --- | --- | --- |
| Trait 1 | Trait 2 | Hypothesis | No. of species | Assumptions on the root | r | HPD r | ESS |
| Male gamete size (continuous) | Separate sexes/co-sexuals | Separate sexes may tend to have smaller and more abundant gametes to ensure reproduction | 38 | NA | 0.0908 | -0.3670, 0.5566 | 525 |
| Male gamete size (continuous) | Male parthenogenesis capacity | There is a minimum male gamete size where parthenogenesis is not possible | 26 | NA | **0.4242\*\*** | -0.0616, 0.8494 | 390 |
| *Two discrete binary traits* |  |  |  |  |  |  |  |
| Trait 1 | Trait 2 | Hypothesis | No. of species | Assumptions on the root | InL Indep | InL Dep | Log BF* |
| Separate sexes/co-sexuals | Parthenogenesis in female | Separate sexes may promote parthenogenesis as a mechanism to ensure reproduction | 40 | 1,0 = Sexes on separate thallus, no parthenogenesis in female | -38.9826 | -40.8219 | **-3.6787** |
| Sexual system (only monoicous and dioicous taxa) | Life cycle dominance | Monoicous species are expected to have haploid dominant phase | 43 | 1,0 and 1,1 = Dioicous, but for the life cycle dominance both states could be the root. | -50.7461 | -48.9906 | **3.5110** |
| Level of gamy (isogamous versus anisogamous) | Life cycle dominance | Anisogamous species are expected to have dominant diploid stages | 68 | 1,1=anisogamy, diploid dominant | -50.7400 | -50.7940 | -0.1080 |

\*BF > 2 is considered evidence of the dependent model
ESS: effective sample size
HPD: highest posterior density
\*\*significant different from zero according to the threshold model

**Table 4.** Estimated rates of correlated transitions

| Rate | State from | State to | Sexual system and<br>life cycle dominance |  |
| --- | --- | --- | --- | --- |
|  |  |  | Mean | sd |
| q12 | 0,0 | 0,1 | 0.0118 | 0.1683 |
| q13 | 0,0 | 1,0 | 0.0153 | 0.1682 |
| q21 | 0,1 | 0,0 | 0.0080 | 0.0090 |
| q24 | 0,1 | 1,1 | 0.0160 | 0.1682 |
| q31 | 1,0 | 0,0 | 0.0094 | 0.1683 |
| q34 | 1,0 | 1,1 | 0.0036 | 0.1681 |
| q42 | 1,1 | 0,1 | 0.0105 | 0.1682 |
| q43 | 1,1 | 1,0 | 0.0034 | 0.1681 |

**Figure 4.**
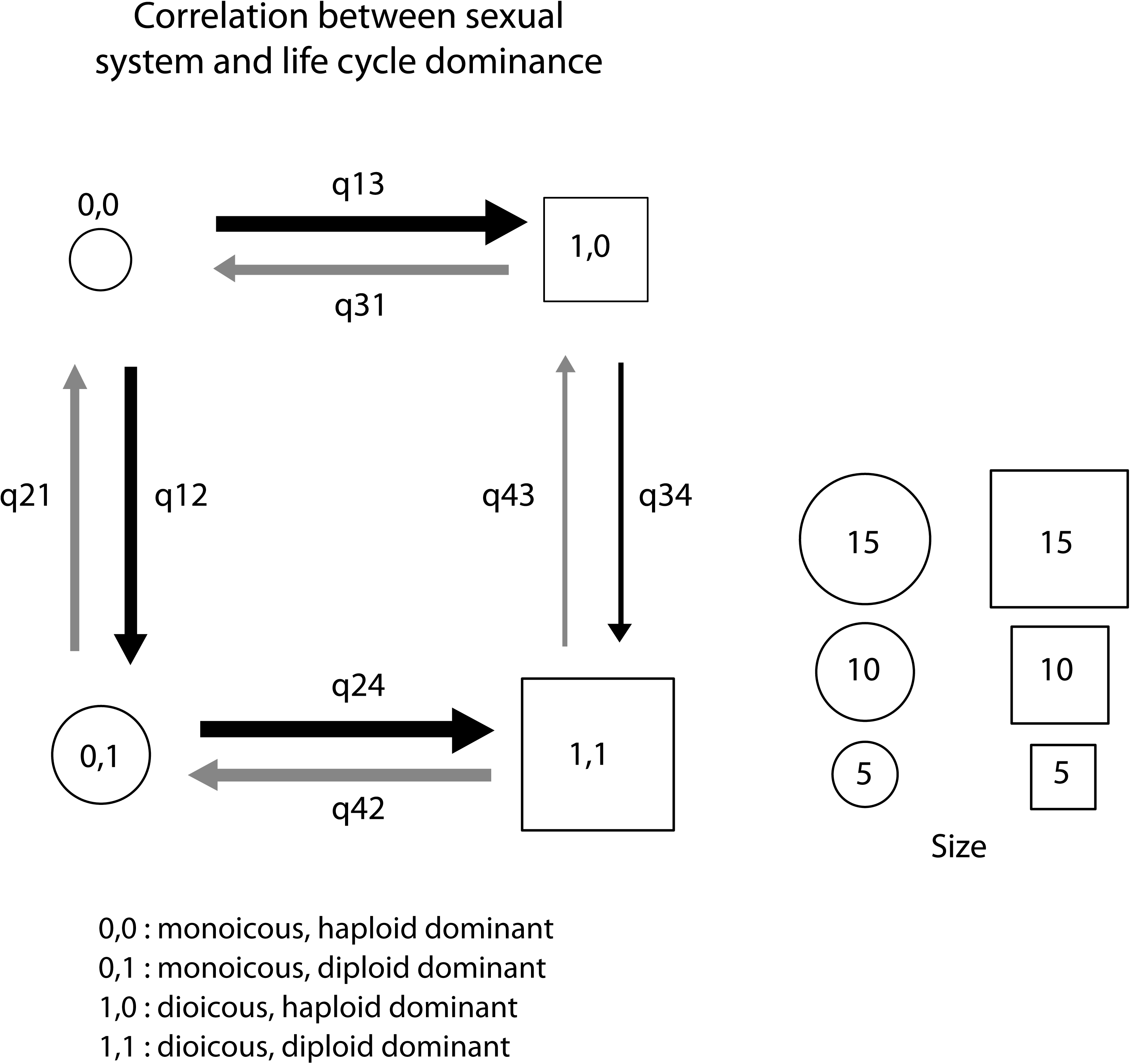
Models of the correlation between traits in brown algae. Binary states are described at the bottom of each panel. Transition rates follow the nomenclature in BayesTraits software (http://www.evolution.rdg.ac.uk/BayesTraitsV3.0.1/BayesTraitsV3.0.1.html). Square boxes indicate the most likely ancestral state (see ancestral state estimations in Methods), sizes of squares/circles represent the abundance of each state in the set of sampled species, and arrow thickness is proportional to the rate of transition between states (see Table 2).

### Generation dominance and sexual dimorphism

We tested if diploid dominance is correlated with an increase in sexual dimorphism. The test of the dependent versus independent model showed that the difference in likelihood was not significant (log BF = −0.1080, Table 3), suggesting that the evolution of the traits is not correlated. Therefore, our data does not support the hypothesis that diploid growth is associated with increased sexual dimorphism.

### Gamete biology and sexual systems

Based on the idea that gamete dimorphism evolved to maximize the chances of gamete encounters, one may hypothesise that separate sexes (dioecy and dioicy) would be associated with small and abundant male gametes, as a mechanism to ensure that the gametes find a partner of the opposite sex when gametes are release into seawater. However, we found no evidence for an association between male gamete size and sexual system (sexes on same versus different individuals) (Figure S2, Table 3, r=0.0909).

When gametes are produced by two different individuals, it may be more difficult for a gamete to find a gamete of the opposite sex than if the same individual produces gametes of both sexes. Accordingly, we hypothesised that parthenogenesis would be favoured in species with separate sexes, as opposed to the situation where male and female gametes are produced by the same individual (note that auto-incompatibility has not been described in the brown algae, with the exception of one study, Gibson 1994). However, we found no evidence that parthenogenesis was more prevalent in species with separate sexes (Table 3).

Finally, we investigated the relationship between the size of male gametes and their parthenogenetic capacity, under the hypothesis that there is a minimum threshold size for male gametes, below which parthenogenesis is not possible. The phylogenetic threshold model indicated that there is a positive correlation between male gamete size and parthenogenetic capacity (Table 3, r=0.4242), however the highest posterior density (HPD) interval of this correlation includes zero. We therefore complemented this analysis using an Ornstein Uhlenbeck (OU) model (Hansen 1997; Butler and King 2004; Harmon et al. 2010). This analysis concluded that the estimated optimal size for non-parthenogenetic male gametes is significantly lower than that of parthenogenetic male gametes (5.49 um vs 9.30 um; Figure S1, Figure 5). This analysis therefore highlights a significant association between male gamete size and parthenogenetic capacity.

**Figure 5.**
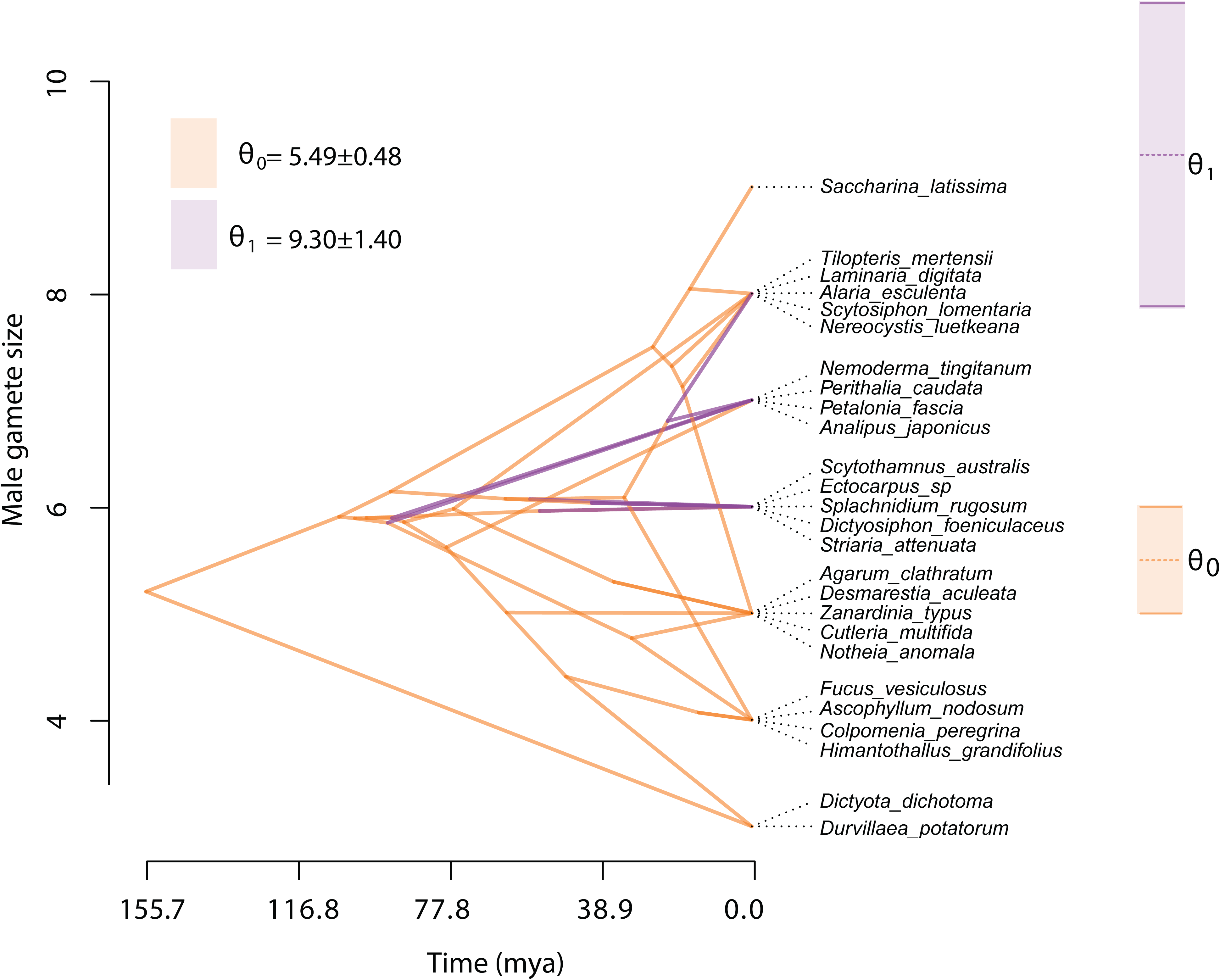
Phenogram for male gamete size and parameter estimation using Ornstein-Uhlenbeck (OU) model. Colours denote the absence (θ0 in orange) and presence (θ1 in purple) of parthenogenesis of male gametes.

## DISCUSSION

### Diploid growth evolved to complement deleterious mutations

Based on the analysis carried out here, the ancestral brown alga appears to have had a diplohaplontic life cycle with similar diploid and haploid dominance (i.e., similar size and complexity of the gametophyte and sporophyte generations). Over evolutionary time, the diploid phase became dominant in some clades (and haploid dominance decreased) whereas other clades evolved towards greater haploid dominance. Several theories have been proposed to explain evolution towards either a dominant haploid or a dominant diploid phase in the life cycle (e.g., Otto & Gerstein 2008). Hypotheses based on the effect of deleterious alleles have proposed that being diploid generally increases mean fitness due to the masking of deleterious alleles (due to complementation of these alleles by non-mutant alleles), while developing as a haploid allows more efficient purging of deleterious alleles because they are exposed to selection (Otto & Goldstein 1992, Rescan et al 2016, Scott & Rescan 2017). The balance between these two forces determines whether evolution proceeds towards an increase of the haploid or the diploid phase, and depends critically on the importance of sexual exchanges within populations. Indeed, under higher rates of inbreeding or asexual reproduction, the benefit of purging deleterious alleles remains associated with alleles increasing the haploid phase, therefore haploidy is favoured. In contrast, outcrossing and/or more frequent sex tend to favour diploidy (Otto & Marks 1996). Very few estimates of inbreeding coefficients or rates of asexual reproduction are available for brown algae but this idea was tested by Bell (1997) by looking at the correlation between the sexual system of a species (monoicous or dioicous) and the relative dominance (i.e., size) of the haploid and diploid phases of the life cycle, assuming that monoicous species will tend to be more inbred due to selfing. At the time, Bell concluded that monoicous species did not tend to have more dominant haploid phases (Bell 1997). In contrast to Bell’s analysis, which was based on a small number of brown algal species, our results do appear to support Otto and Marks’ ideas, at least to some extent, because transitions towards dominance of the haploid phase were found to be more frequent when the sexual system was monoicous, consistent with the idea that monoicy is correlated with haploid growth. Estimates of inbreeding coefficients within natural populations of monoicous species would be extremely valuable to shed further light into these phenomena.

Somatic mutations have been proposed as another possible source of selection for diploidy, as these mutations should have a lower impact on the fitness of diploid organisms (e.g., Otto & Gerstein 2008). This idea is consistent with the general observation that larger organisms tend to be diploid rather than haploid. Indeed, this pattern holds true for the brown algae, since all the largest brown algae (e.g., Laminariaceae, Sargassaceae, Fucaceae) have a dominant diploid phase.

Bell (1997) has also proposed that the different biology of spores and gametes should select for larger sporophytes (allowing efficient dispersal of spores) and smaller gametophytes (so that gametes are released close to the substratum, in order to facilitate gamete encounters). While this type of constraint may explain why many brown algal species have a smaller gametophyte, it does not explain the evolution towards larger gametophytes that occurred in some clades, nor the transitions towards diploid cycles in which gametes are released by large diploid individuals.

### Diploid growth is associated with increased gamete dimorphism

The theory for the evolution of gamete dimorphism based on the trade-off between gamete number and offspring fitness predicts that dimorphism may evolve when zygote size has a strong effect on fitness (i.e. when offspring fitness increases more than linearly in relation to zygote size). Accordingly, one may predict that if a larger zygote size is needed for larger diploid development, increased diploid growth would favour higher levels of gamete dimorphism (Parker et al. 1972). Bell (1994) proposed an alternative theory that also predicts a correlation between diploid growth and gamete dimorphism, in which the direction of causality is reversed, i.e., sexual selection caused by gamete dimorphism favouring diplontic cycles in order to increase genetic differences between gametes produced by the same organism. Although our results show that diplontic brown algal species are exclusively oogamous, suggesting a link between strong sexual dimorphism and diploidy, those associations may not be relevant because they are based on only one group, the Fucales. Therefore, overall, our analyses have not allowed to validate the idea that diploid growth is associated with increased sexual dimorphism.

### Evolution of sexual systems in the brown algae

The ancestral sexual system of brown algae corresponds to haploid sex determination and dioicy, with several transitions towards monoicy having occurred independently over evolutionary time. Transition towards a diplontic life cycle in the Fucales involved a monoecious/hermaphrodite intermediate state, with subsequent independent re-emergence of dioecy in some lineages. It is interesting to note that transitions from separate sexes to co-sexuality are relatively frequent in haploid sexual systems, which contrasts to what is the most commonly accepted direction of evolution in clades with diploid sex determination (i.e., monoecy to dioecy). Note, however, that although dioecy was considered to be an evolutionary dead end in angiosperms, more recent phylogenetic analysis are challenging this conclusion, and the idea that reversals to monoecy in angiosperms may be more frequent than thought before is increasingly becoming accepted (Kafer et al. 2017; Pannell 2017).

In diploid sexual systems, two main selective effects have been proposed to explain transitions from co-sexuality to dioecy: inbreeding depression (selfing is less likely to occur when male and female gametes are produced by separate individuals) and the effect of trade-offs between male and female fitness (Charlesworth & Charlesworth 1978, Charnov 1982). In haploid sexual systems, the opposite transition (from separate sexes towards co-sexuality) could also in principle be caused by a change in the shape of the trade-off between male and female reproductive success (leading to a higher fitness of gametophytes producing both types of gametes) or by selection for inbreeding (selfing), either through the automatic transmission advantage associated with selfing (Fisher 1941) or for reproductive assurance when population density is low. Note, however, that parthenogenesis occurs in all monoicous species, and this process may represent an alternative way of dealing with mate limitation and reproductive assurance. Assuming that selfing occurs following transitions to monoicy, such transitions should occur more easily when inbreeding depression is low. One would therefore expect more transitions to monoicy in taxa with a prolonged haploid phase (assuming that at least a proportion of the deleterious alleles affecting the fitness of diploids will be purged during the haploid phase of the life cycle), but this is not what we observe here (q31<q42, Fig. 4). Examining the proximate mechanisms involved in the transitions between separate sexes and co-sexuality in both haploid and diploid systems and more natural population data, for example in populations with different densities, would be valuable to shed light on the mechanisms and evolutionary forces driving the shifts among sexual systems in the brown algae.

### Anisogamy is ancestral in the brown algae

In agreement with the conclusion obtained by Silberfeld et al (2010), our analysis points towards an oogamous ancestor of brown algae, with several independent transitions towards anisogamous and isogamous clades. This stands in contrasts with theoretical scenarios representing the evolution of gamete dimorphism from an isogamous ancestor (e.g., Sanderson & Hurst 2001, Parker et al 1972; Charlesworth 1978, Lehtonen and Koko 2011; Lehtonen et al 2016), in which isogamy is ancestral and anisogamy represents an intermediate step during the process of increased gametic differentiation. To our knowledge, the possible selective forces favouring reduced gamete dimorphism have not been explored by theoretical models. Theories based on disruptive selection caused by a trade-off between the number of gametes produced and zygote size (e.g., Parker et al 1972, Bulmer & Parker 2002) have shown that the shape of the relation between zygote size and fitness is critical for the evolution of gamete dimorphism. It would be interesting to explore whether a change in the relation between zygote size and fitness (for example, due to a decrease in size of the diploid organism) may favour transitions from oogamy to anisogamy or isogamy. This type of evolutionary mechanism may generate a positive correlation between the degree of gamete dimorphism and the relative importance of the diploid phase, leading to an inversion of the causal relationship in Bell’s (1994, 1997) hypothesis mentioned above, i.e., decrease in the size of the diploid organism would drive a decrease in gamete dimorphism.

Lehtonen and Kokko showed that evolution of anisogamy requires some level of gametic competition and limitation (Lehtonen and Kokko 2011). Therefore, it is likely that in specific conditions the system may return to isogamy or near isogamy, for instance if there is a low level of gamete competition or if there is no gamete limitation.

### Evolution of gamete size and parthenogenetic capacity

There are marked differences between the relative parthenogenetic capacities of male and female gametes in isogamous, anisogamous and oogamous brown algal species (Luthringer et al. 2015). Overall, both male and female gametes of isogamous species are capable of parthenogenesis, whereas only the female gametes of anisogamous species are parthenogenetic. In many oogamous species neither the male nor the female gametes undergo parthenogenesis. It has been suggested that increased gamete size leads to increased parthenogenetic capacity, up to a point, but that in oogamous species, the large female gamete loses its flagella becoming specialised for zygote production and is no longer capable of initiating parthenogenetic development (Luthringer et al. 2015).

The gametes of the ancestral brown alga seem to have been unable to perform asexual reproduction through parthenogenesis, suggesting that the emergence of gamete parthenogenetic capacity was a derived, and perhaps adaptive trait for instance in marginal populations or other situations where mates are limited (Stalker 1956; Bierzychudek 1987; Oppliger et al. 2014). Data from field populations of a range of species would be needed to further understand whether parthenogenesis is adaptive. It is noteworthy that parthenogenetic capacity is assessed under laboratory conditions, and that the contribution of parthenogenesis to recruitment in natural populations would be worth exploring further (Oppliger et al. 2007, 2014). Note that a recent study in field populations of the Ectocarpus showed no evidence that parthenogenesis plays a significant role under field conditions (Couceiro et al. 2015). In contrast, studies in field populations of another brown algal species, *S. lomentaria* suggested that parthenogenesis is prevalent in field populations (Hoshino et al. 2018). Interestingly, female-only parthenogenetic populations have larger gamete sizes relative to ‘sexual’ populations of the same species, consistent with a link between gamete size and parthenogenetic capacity, and opening the possibility that parthenogenesis may be an adaptive trait.

## Methods

### Molecular data

Alignments were based on the data published by Silberfeld and colleagues (Silberfeld et al. 2010) that included five mitochondrial genes (*atp9:* mitochondrial ATP synthase subunit 9 gene, *cox1* and *cox4:* Cytochrome c oxidase subunit 1 and 3 genes, *nad1* and *nad4:* NADH dehydrogenase subunit 1 and 4), four plastid genes (*rbcL:* large subunit of plastid encoded ribulose-1,5-biphosphate carboxylase oxygenase gene, *psaA:* photosystem I P700 chlorophyll a apoprotein A1 gene, *psbA:* photosystem II protein D1 gene, and *atpB:* ATP synthase subunit b gene), and a nuclear gene (*LSU:* large subunit of 28S rRNA gene). Our final tree contained 131 species. To attribute trait states to the species in this tree we replaced some entities, depending on the availability of life-history information (i.e., kept the sequence data used to build the tree but used the data on life-history from another close relative) and added sequences from Genebank for species of the genera *Padina, Sargassum, Alaria* and *Ectocarpus* (Table S1). No information was available about the life histories of the closest relatives of the Phaeophyceae, e.g. Phaeothamniophyceae, so we used *Vaucheria*, a siphonous genus in the heterokont class Xanthophyceae for which life cycle and reproductive trait information are available, as an outgroup. The final species list used for the trait analysis, for which we had life cycle and reproductive trait information, was comprised of 77 species, including the outgroup.

### Phylogenetic reconstruction

All sequences were aligned using MAFFT (Katoh et al. 2009), and the best substitution model was estimated using the phymltest function in the *ape* R package (Paradis et al. 2004). The concatenated alignment was used for Bayesian Inference with Beast v1.8.2 (Drummond et al. 2012) with three different gene partitions, for nuclear, plastid and mitochondrial genes. Each partition was unlinked for the substitution model. We used birth-death with incomplete sampling as tree prior, and four calibration nodes as described in (Silberfeld et al. 2010) (see nodes A to D, Figure 2). We used log-normal priors for two of the calibrations: *Padina*-like clade A, lognormal distribution (mean 5 Ma, sd 1, and lower boundary at 99.6 Ma); *Nereocystis-Pelagophycus* clade B: lognormal distribution (mean 20 Ma, sd 1, and lower boundary at 13 Ma), and normal priors for the root (Phaeophyceae root age D: normal distribution (u=155, sd=30 Ma), and the Sargassaceae node C to a normal distribution (u=60, sd=15, with lower boundary 13 Ma). Finally, the MCMC was set to 50 million generations with a sampling every 1,000. The posterior distribution was summarized using Treeannotator v1.7.0 (Drummond et al. 2012) to obtain a Common Ancestor Tree (Heled and Bouckaert 2013). For the macroevolutionary analyses (see below), a set of 100 trees were sampled from the posterior distribution.

### Life history traits

Four life-history traits were recorded based on a literature review: life cycle type (haploid > diploid; haploid = diploid; haploid < diploid; diplont), sexual system (monoicous; dioicous; monoecious; dioecious), gamete dimorphism (isogamous; anisogamous; oogamous), and the occurrence of gamete parthenogenesis (no parthenogenesis; parthenogenesis in female gametes only; parthenogenesis in both male and female gametes). The traits were coded as discrete multi-state characters. A full explanation of each state is given in Table S2. We separated the respective traits into seven additional characters. For example, we transferred ‘gamete size’ (iso-, aniso-, oogamous) into a continuous male gamete size trait. We furthermore recoded multi-state traits into binary data for the correlation tests (see below), such as the ‘gamete dimorphism’, which was recorded by separating the absence (0 = oogamy) from presence (1 = iso-or anisogamy) of female flagellated gametes. We categorized an additional sexual system trait as ‘sexes occurring on the same thallus’ (0 = monoicous or monoecious) or ‘separate thalli’ (1 = dioicous or dioecious). The life cycle was simplified to the occurrence of a ‘dominant haploid phase’ (0 = haploid ≥ diploid) versus dominance of the diploid phase (1 = haploid < diploid or diplontic), with dominance broadly meaning size of the adult individual. Finally, the occurrence of parthenogenesis was separated into two additional traits, absence (0) or presence (1) of male parthenogenesis, and absence of parthenogenesis (0) versus parthenogenesis occurring in at least one of the sexes (1), most commonly the female.

For simplicity, we coded as “isogamous” algae with physiological and behavioural anisogamy but that have been described as having no size difference between male and female gametes. Note that all brown algae exhibit an asymmetry between male and female, at least at the level of their behaviour, and potentially all the algae scored as isogamous have in fact subtle size differences but the literature is not detailed enough in this respect. For example, most representatives of the order Ectocarpales have been reported to be ‘isogamous’ (based on observations under the microscope, but without detailed measurements of gamete size), but some members (*Ectocarpus* sp., *Colpomenia peregrina* Sauvageau) as well as the last species branching off before the Ectocarpales, *Asterocladon interjectum*, have anisogamous male and female gametes.

The sister group to the Ectocarpales, the order Laminariales, is almost completely oogamous, with the exception of the genus *Saccharina*, which has been shown to have eggs with rudimentary flagella (Motomura and Sakai 2008) being therefore considered strongly anisogamous.

### Ancestral state reconstructions

A likelihood-based method was used to reconstruct the ancestral state of each of the four life-history traits. We fitted three different models of trait evolution using the function *fitDiscrete* from the R package Geiger (Harmon et al. 2008). These models differed in the number of transition rates as follows: *equal rates* (ER, a single transition rates between all states), *symmetric* (SYM, forward and reverse transitions are the same), and *all-rates-different* (ARD, each rate is a unique parameter). The corrected Akaike Information Criterion (AICc) was used to compare the alternative models. Each model was estimated on each 100 phylogenetic trees sampled from the posterior distribution to account for uncertainty in tree topology and divergence times. State probabilities at the root and transition rates were summarized with the mean and standard deviation values of all iterations, to incorporate phylogenetic uncertainty.

We inferred the number of transitions between states, and their minimum timing, using stochastic character mapping (Huelsenbeck et al. 2003). One hundred stochastic mappings were performed on the posterior sample of trees, and on each we divided branch lengths into time bins of 1 Myr and recorded the number of transitions from and to each state, in each bin (as described in (Serrano-Serrano et al. 2017). We reported the mean and standard deviation, and the time bin at which 60% of the stochastic mappings had at least one transition event as the onset time for each type of transition.

### Correlation analyses

We first assessed correlation between life history traits using a reversible-jump MCMC algorithm to test the correlation between two binary traits as implemented in *BayesTraits* V3 (Pagel *et al.* 2004). This approach compared two models, a null model assuming that the traits had evolved independently, and an alternative model assuming that their evolution had been correlated. Each model was run for 10 million generations using the values found in the ancestral state reconstructions for the root state. The two models were compared through their log marginal likelihood by estimating the log Bayes factor. This approach was used to test the correlation between female parthenogenesis and the occurrence of sexes on the same versus separate thalli. Tests showing a significant support for the correlated model were presented as networks of evolutionary transitions using the R package qgraph (Epskamp *et al*. 2012). Second, we used the implementation of the threshold model, *threshBayes* in the R package phytools (Revell 2014), to test for the correlation between a continuous and a discrete variable. The threshold model assumes that the states of discrete phenotype are governed by an unobserved continuous character called *liability.* These liabilities are assumed to evolve according to a Brownian motion model (Felsenstein 2012) and translate into discrete characters once they have passed certain thresholds. We used this model to test the correlation between male gamete size and two discrete traits, male parthenogenesis and sexes on the same or on separate thalli. For correlation analyses that were significant, we fitted an Ornstein–Uhlenbeck model of evolution to further test whether the continuous trait had two discrete selective regimes, determined by the discrete binary trait. We used the *OUwie* from the R package Ouwie (Beaulieu *et al.* 2012), and compared the alternative models using the corrected Akaike Information Criterion (AICc)-selected model.

## SUPPLEMENTAL FIGURES

**Figure S1.** Phylogenetic tree pruned to the 43 species with sexual system and life cycle dominance traits recorded. Coloured squares at the tips represent the species state for sexual system (left column) and phase dominance (right column). Colour codes for each binary trait are explained in the top left legend. Ancestral node states are not depicted, see Figure 3 for full ancestral state reconstruction.

**Figure S2.** Evolutionary correlation between continuous and discrete traits using the threshold model. **A,** Correlation between the size of male gametes and presence of sexes on same/different individual (unisexual versus co-sexuals). **B,** Correlation between the size of male gametes and male parthenogenesis. Blue histograms (left panel) denotes the posterior density for the correlation between traits. Posterior liabilities, or unobserved continuous traits, for each discrete trait (right panel). HPD: highest posterior density.

## SUPPLEMENTAL TABLES

**Table S1:** Accession number for the sequences of the species that were not included in (Silberfeld et al. 2010).

**Table S2:** List of detailed life cycle and reproductive traits across the brown algal species

## SUPPLEMENTAL DATASET

Details of all the traits and species used in this work, including references.

