## Supplementary figures and images for "Evolution of life cycles and reproductive traits: insights from the brown algae"

### Figure S1

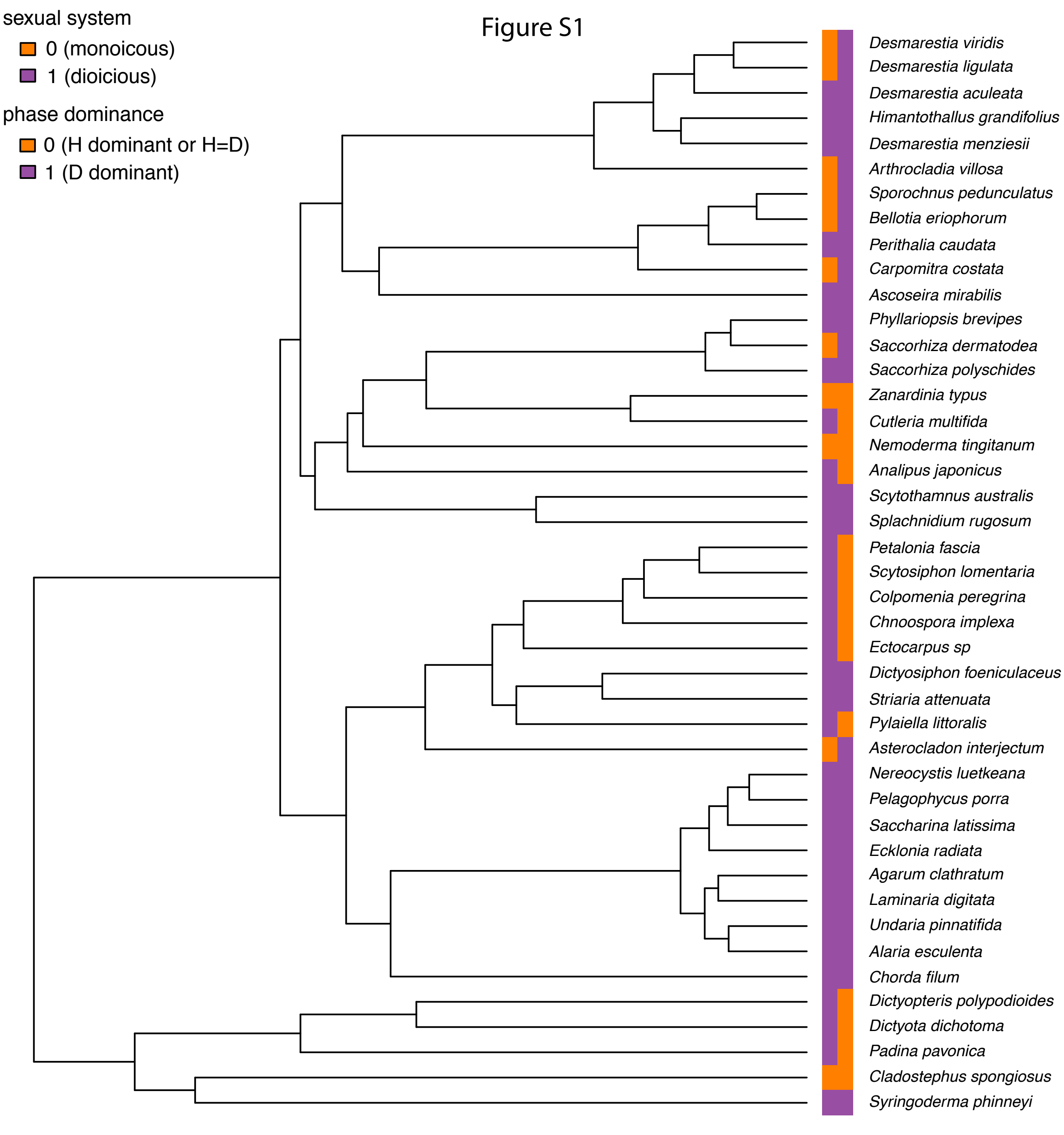
