## Supplementary material for "Evolution of life cycles and reproductive traits: insights from the brown algae": Figure S2

Correlation (R) between male gamete size (continuous) and sexes on thallus (discrete)

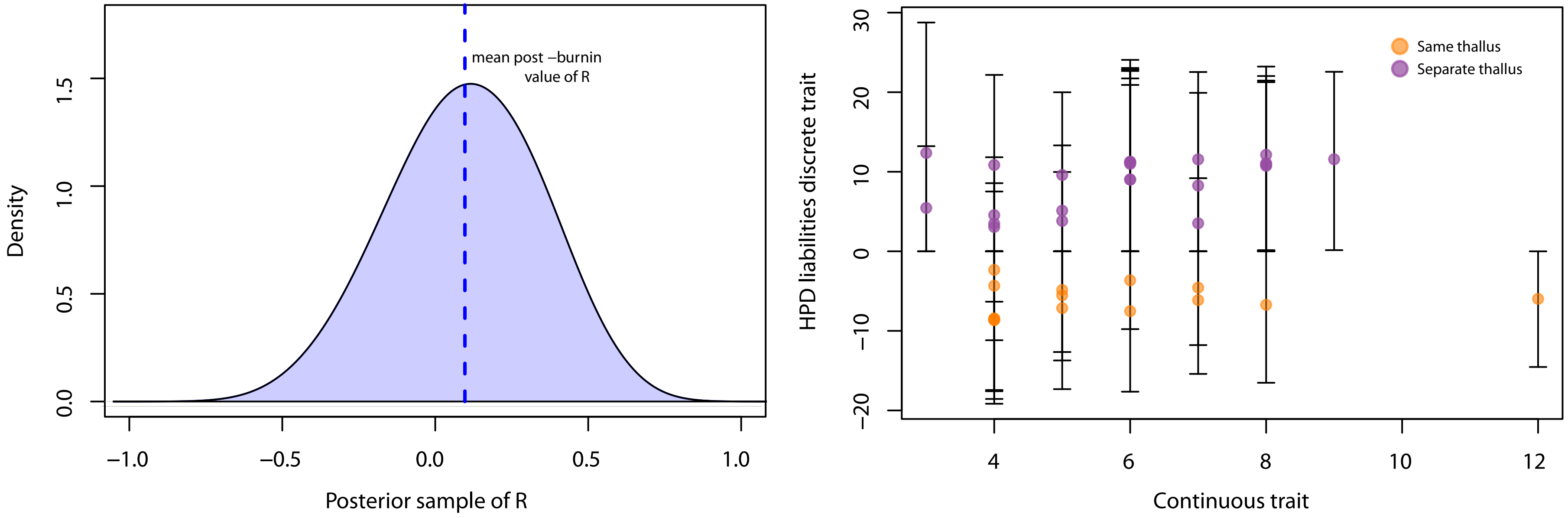


Correlation (R) of male gamete size (continuous) and male partenogenesis (discrete)

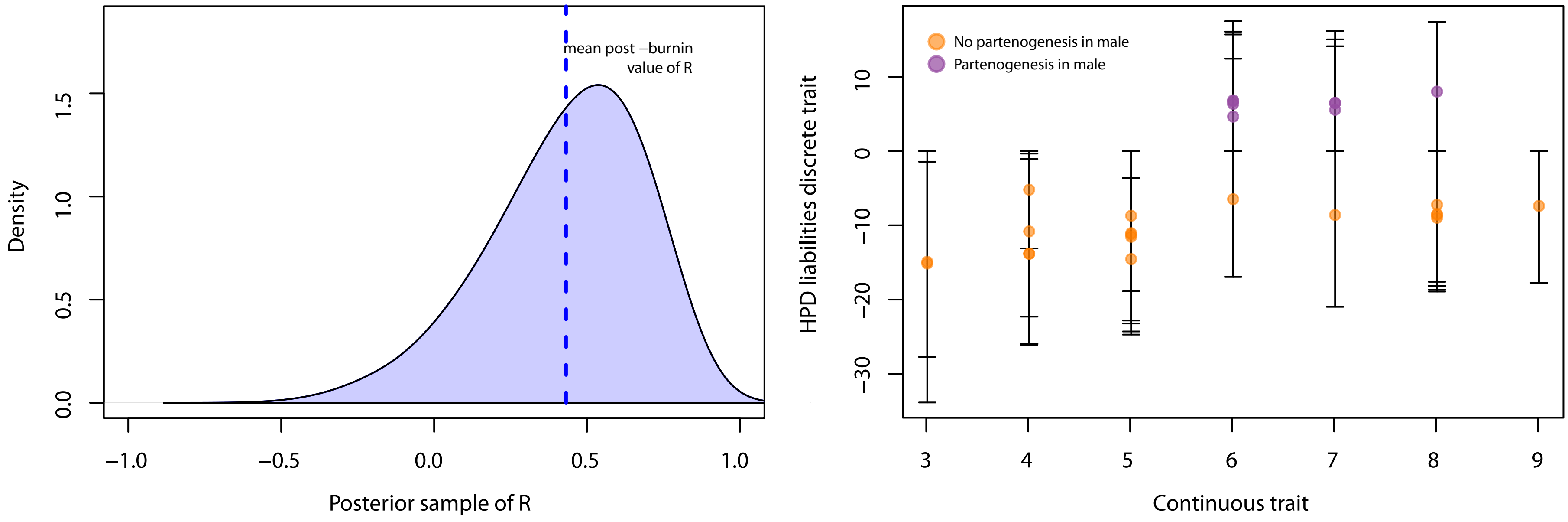
